# The sum and the parts: dynamics of multiple and individual metastases during adaptive therapy

**DOI:** 10.1101/2022.08.04.502852

**Authors:** Jill Gallaher, Maximilian Strobl, Jeffrey West, Jingsong Zhang, Robert Gatenby, Mark Robertson-Tessi, Alexander R. A. Anderson

**Affiliations:** Department of Integrated Mathematical Oncology, Moffitt Cancer Center; Department of Genitourinary Oncology, Moffitt Cancer Center; Department of Radiology, Moffitt Cancer Center

## Abstract

Evolutionary therapies, such as adaptive therapy, have shown promise in delaying treatment resistance in late-stage cancers. By alternating between drug applications and drug-free vacations, competition between sensitive and resistant cells can be exploited to maximize the time to progression. However, the optimal schedule of this dosing regimen depends on the properties of the tumor, which often are not directly measurable in clinical practice. In this work, we propose that the initial cycle of adaptive therapy can be used as a tool to probe the relevant tumor properties. We present a framework for estimating individual and collective components of a metastatic system through tumor response dynamics, which uses a system of off-lattice agent-based models to represent individual metastatic lesions within independent domains, all of which are subject to the same systemic therapy. We find that the first cycle of adaptive therapy delineates several features of the computational metastatic system: larger metastases have longer cycles; a higher proportion of drug resistant cells slows the cycles; and a faster cell turnover rate speeds up drug response time and slows the regrowth time. The number of metastases does not affect cycle times, as response dynamics are dominated by the largest tumors rather than the aggregate. In addition, the heterogeneity of the system is also a guide for therapeutic approaches: generally, systems with more between-tumor (intertumor) heterogeneity had better success with continuous therapy, while systems with more within-tumor (intratumor) heterogeneity responded better to adaptive therapy. Intertumor heterogeneity was found to correlate more with dynamics from patients with high and low Gleason scores while intratumor heterogeneity was correlated with dynamics from patients with intermediate Gleason scores.

## Introduction

The majority of cancer-related morbidity and mortality is driven by late-stage metastatic disease [1]. Metastasis puts an increasing metabolic strain on the body and disrupts normal tissue function, which causes an overall weakening of the patient. Systemic cytotoxic treatments can lessen the overall disease burden, but a balance is needed between maximizing tumor kill, avoiding patient toxicity and poor quality of life, and managing the emergence of treatment resistance [2]. Whilst often metastatic staging may be binary (metastases present or not), some cancers have more nuanced classifications, such as the oligometastatic state, where metastases-directed treatment may be curative [3]. Treatment failure occurs due to resistant cells, which can arise from all lesions, from a subset of them, or from a single metastatic site. Given the wide variation in lesion sizes, locations, and characteristics, measuring the relevant properties of metastatic disease that can inform treatment decisions is inherently challenging. There is potential heterogeneity in the molecular make-up of a single metastatic lesion as well as between lesions. Whilst large tumors can be routinely monitored by imaging, newly colonized micrometastases may be too small to be detected. In some cancers, overall burden and disease state may be estimated with blood biomarkers, but at best this provides a coarse-grained view of systemic burden. Thus, new ways in which to measure features of metastases and inter- and intra-lesion heterogeneity of metastatic cancers are urgently required.

In this work, we focus on metastatic prostate cancer. The clinical data that is routinely collected consists of blood biomarkers, imaging, and biopsies. Prostate specific antigen (PSA) is a blood biomarker collected from peripheral blood, which is routinely used for prostate cancer screening. PSA is produced by most prostate cancer cells and is the gold-standard indicator for systemic tumor burden. Diagnostic imaging with either a bone scan, local MRI, or CT can be used to visualize the extent of metastases that most often seed to the bone, and less often to the lymph, lung, and liver. Biopsies of the primary lesion yield a Gleason score, which grades the level of similarity between benign and tumor tissue. Immunohistochemistry and other assays may be used to identify various markers in the primary lesion and potentially some larger metastatic lesions. However, each of these sources of data has its drawbacks. Biopsies can only capture a single timepoint for a small section of a visible and accessible lesion. Imaging, while able to capture whole lesions over multiple time points, is more spatially coarse-grained and has a threshold of detection that misses small lesions. Furthermore, bone scans for metastatic prostate cancer often capture metabolically active hotspots that can indicate the presence of a lesion, but quantifying its size is obscured by inflammation. Finally, blood biomarkers such as PSA, circulating tumor cells, and circulating tumor DNA can provide a finer temporal resolution, but only represent the total systemic burden, if not just a subset of the total burden.

PSA is the most widely used molecular marker for the diagnosis and management of a malignancy, however, absolute PSA alone is not always useful [4]. Prior to therapy, it is not clear whether PSA is prognostic, however, PSA ≤ 4 ng/mL after 7 months of androgen deprivation therapy strongly predicts improved survival [5]. While most studies have focused on using static metrics to predict outcome, an emerging approach which holds promise is using dynamic biomarkers instead. These markers base prognosis not on the absolute value of measurement at a single time point, but on the relative change over time. For example, PSA doubling time may stratify prostate cancer patients likely to respond better to chemotherapy, progress to metastatic disease, or die from prostate cancer [4]. In this paper we explore how PSA response dynamics can allow us to infer additional information about the characteristics of the disease that standard analysis of biomarkers cannot reveal.

To do so, we leverage data from a new treatment approach which has been gaining ground in the treatment of advanced cancers. This is an evolutionary-informed strategy, termed “adaptive therapy”, in which dosing is adjusted in response to biomarker dynamics. The aim is to prolong control over the disease by maintaining a certain tumor burden in exchange for delaying, or even preventing, the emergence of treatment resistance. Treatment breaks are given adaptively in a patient-specific manner, in order to prevent “competitive release”, which is when a small population of resistant cells begins to grow after therapy has removed the larger population of sensitive cells that was competitively holding resistant growth in check. Adaptive therapy has been successfully tested in several *in vivo* models [6]–[8], which has motivated the launch of clinical trials in prostate cancer, thyroid cancer, and melanoma (NCT02415621, NCT03511196, NCT03630120, NCT03543969). The most advanced of these clinical studies investigated an adaptive algorithm for the treatment of metastatic castrate-resistant prostate cancer (mCRPC) using the Cyp 17 inhibitor abiraterone [9]. In this study, tumor burden was monitored monthly using PSA, and treatment was withheld once patients had achieved a 50% drop from their pre-treatment PSA level. There have been other trials that have used similar dose-minimizing algorithms, such as intermittent therapy [10]–[13]. However, these studies started and stopped therapy based on either fixed time intervals, or absolute values of PSA as a threshold. In contrast, the adaptive therapy study normalized the PSA thresholds for each patient by their initial PSA levels; this allows adaptive therapy to be more sensitive to the nadir of cancer burden in each patient, which must remain high enough to prevent competitive release of resistant cells. In addition, this patient-specific approach may reveal additional insights into the tumor’s dynamics when coupled with mechanistic mathematical models.

Mathematical modelling has played an important role in advancing the development of adaptive therapy [9], [14]–[26]. These studies have mainly focused on how various features of sensitive and resistant cells affect the response to adaptive therapy, and in general the models applied to individual tumors and systemic biomarkers, rather than widely disseminated metastatic disease. We take a different approach and consider the treatment response dynamics in order to characterize the properties of the system of metastases. Relatively few mathematical models have been explicitly tailored to understanding metastases at all; those that have cover a diversity of aspects pertaining to metastatic disease, which can be represented simply as a burden [3], comprehensively by connecting primary tumor growth to metastatic burden [27]–[32], or be defined by the steps in the metastatic cascade [33], [34]. Other models focus on specific aspects of metastasis, such as invasion and survival of metastatic clusters in the circulation, seeding timing, diversity, and initiation [35]–[38], metastatic spread and dissemination [39], the role of Immune system [40], and secondary seeding [41].

In this work, we propose a framework for estimating individual and collective components of a metastatic system using tumor dynamics during the first adaptive therapy treatment cycle. We present an analysis of the longitudinal PSA dynamics of 16 mCRPC patients undergoing adaptive androgen deprivation treatment (NCT02415621). We quantify on- and off-treatment times and investigate relationships with clinical variables such as Gleason score, the change in the number of metastases over a cycle, and the total number of cycles over the course of treatment. To gain a more mechanistic understanding of how the observed statistical trends may relate to characteristics of the underlying disease, we compare our clinical observations with an agent-based model of a metastatic cancer. We investigate how the mode of drug action, total tumor burden, number of metastases, turnover rate, degree of resistance, and inter- and intra-tumor heterogeneity all have distinct effects on the cycle dynamics under adaptive therapy. Whilst we focus on prostate cancer, our theoretical insights may be applied more broadly. As such we propose that biomarkers based on the dynamics in the first cycle of adaptive therapy may allow us to better characterize an individual patient’s metastatic disease and help us to better identify which patients are best suited to adaptive versus maximum tolerated dose therapies.

## Materials and Methods

### Clinical Data

We utilize the clinical data from the 16 mCRPC patients on an adaptive abiraterone therapy trial (NCT02415621). Patients in this trial were given abiraterone until their PSA dropped to 50% of the original value and there was imaging confirmation of no progression [9]. Abiraterone was commenced again once the PSA returned to the original patient specific value, and then the cycle was repeated. The data used included Gleason Score, monthly PSA measurements, and the number of lesions at several time points. Some analyses could only use a subset of the cohort (n=14) due to missing information for some patients, so the number of patients for each result lists this value. More information on the patient demographics can be found in [9], [42], and [43].

### Computational Model

Extending our previously published model [44], our computational framework simulates a system of metastases in which the metastatic lesions are modeled using multiple off-lattice agent-based models (ABM). Each instance of an ABM, representing an individual lesion, is set in its own domain, but is subject to the same systemic therapy protocol. To limit the complexity, we do not explicitly model the primary tumor but focus just on the metastases and their diversity to investigate how the heterogeneity within and amongst the metastases is affected by the systemic treatment.

#### Basic model setup

Fig. 1 shows an overview of our modeling framework. During the initial growth period, metastases are seeded with various phenotypic compositions. The example in Fig. 1A shows a system of three metastases seeded at different time points. The metastases are allowed to grow until their total burden reaches a preset value; this ends the seeding/growth period and marks the point at which metastases are discovered and the treatment phase begins. The treatment schedule follows the adaptive therapy algorithm used in the clinical trial by Zhang et al [9], where PSA is used to guide treatment decisions, such that treatment is applied until the PSA is 50% of the original value and restarted on returning to the original value. In the computational model, PSA is calculated as the total burden (sum of all cells in all metastases), under the simplifying assumption that all cells equally produce PSA. The burden is monitored at 1-week intervals; this is on a faster time scale than the clinical trial in order to account for the smaller spatial scale used in the models. After the total tumor burden is reduced to the 50% threshold, the drug is discontinued. The metastases are allowed to grow unimpeded until reaching the original burden again, at which point the drug is reapplied. Note that if the tumor burden does not decline below 50% due to increasing resistance, the drug will stay on for the duration. The treatment cycles repeat until progression occurs, which we define as the time at which the tumor burden exceeds 120% of the original burden, or the time at which the simulation was terminated upon reaching an imposed time limit. The treatment burden thresholds and time limits vary, and they are specified in the results.

**Figure 1.**
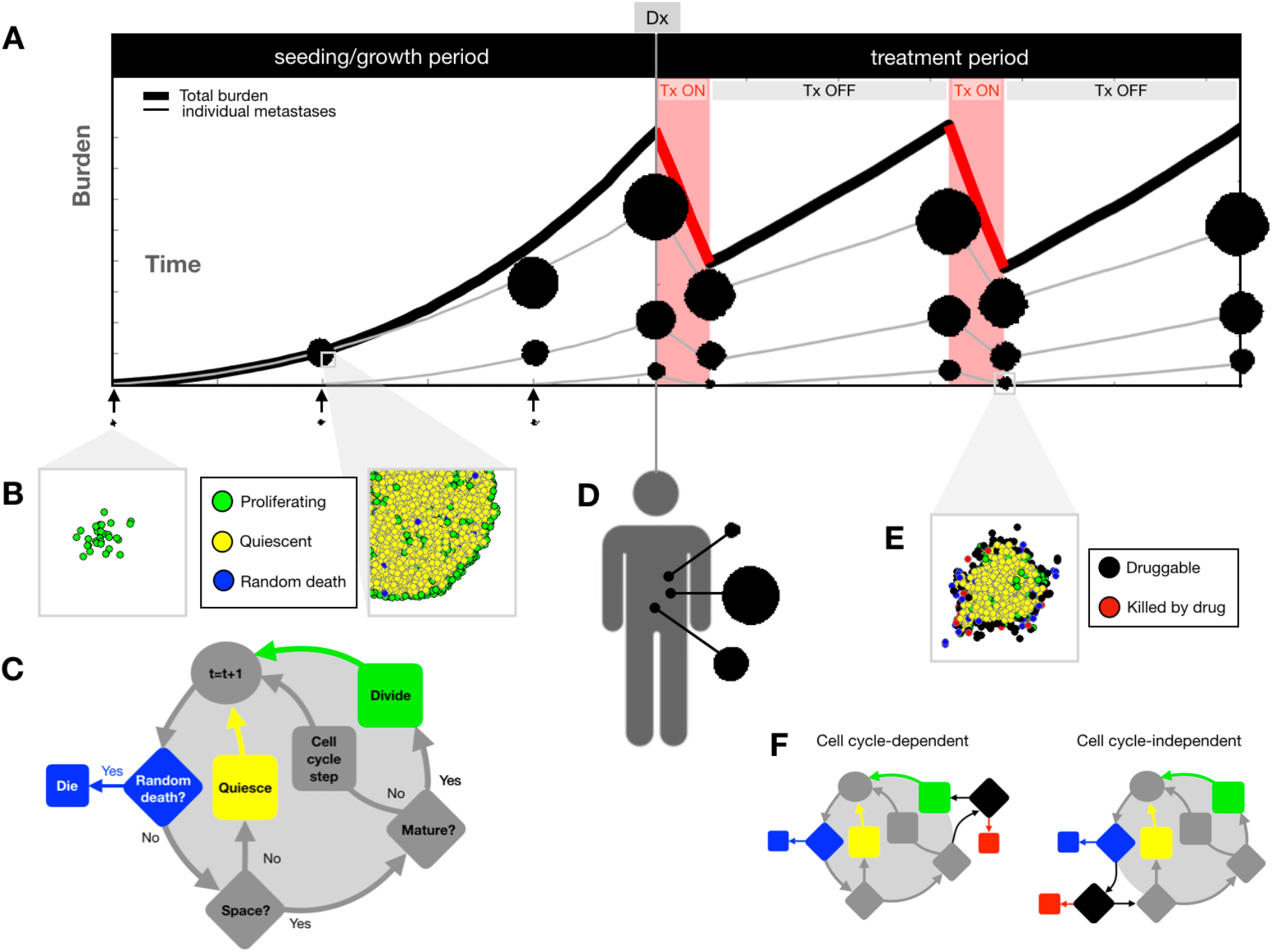
Mathematical model overview and design. A) Simulated treatment trajectory. During the seeding/growth phase, metastases are seeded and grown until reaching a specific total burden at diagnosis (_Dx_). The treatment period follows, during which therapy is applied until the tumor burden reaches half of the original value, and therapy is withdrawn until the tumor burden reaches the original value again, where the treatment cycle begins anew. B) For each metastasis, individual cells are seeded in a separate off-lattice domain. Total burden used for treatment decisions is defined as the sum of all cells in the whole system of metastases. C) ABM decision flowchart for each individual cell over a time step. D) After reaching the designated burden, the treatment period begins. Cells are considered druggable if they lie near the outside edge of the mass (see Supplemental Methods for details), and they can only be killed by the drug if they are druggable. E) Metastases are either subjected to a cell cycle-dependent or cell cycle-independent drug, acting at the point in the decision flowchart shown.

#### Metastasis seeding

Each metastasis is seeded initially with 30 cells. This allows a small initial seed with a large enough distribution of phenotypes to be well-represented when we have intratumor heterogeneity. The metastases are either seeded all at once or at regular intervals *T*_*seed*_ during the seeding/growth period, and we end the growth period at a specific total tumor burden *N*_*Dx*_. If the seeds are staggered, the rate of seeding is determined to ensure that all metastases will be seeded during the growth period before reaching the preset total burden. In this way, we make the loose assumption that the tumors grow roughly linearly at a rate of *k*. This means the final tumor size of each subsequently seeded tumor will be linearly proportional to the time that it has been growing so that each metastasis has a size *ikT*_*seed*_, where *i* is the order index of the metastasis. Taking the final total burden to be equal to the sum of the number of cells in each metastasis, we can then solve for the time interval between each metastasis seeding *as* 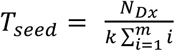, where *m* is the number of metastases, and *k=*150cells/d was calibrated within the relevant parameter space.

#### Cell Interactions and decisions

In the off-lattice agent-based model, which is derived from our earlier work [44], each metastasis is composed of individual tumor cells. These cells vary in proliferation rate, modeled as a rate of progression through the cell cycle. However, proliferation is also subject to each cell’s interactions with its spatial neighborhood (Fig. 1B). The life cycle of a single cell is illustrated in the flow diagram shown in Fig. 1C. At each time point, a cell is subjected to stochastic death (i.e., cellular turnover). If it survives, and if the cell has enough space to proliferate (i.e., there is space to fit a cell adjacent to the dividing cell without any overlap), then it will continue progressing through the cell cycle. If there is not enough space, the cell enters a quiescent state and does not progress through the cell cycle during that time step. If the cell is mature (i.e., it has finished the cell cycle), the cell will divide.

#### Cell Death

Cells may die from either random cell turnover (mentioned above) or from exposure to the drug. Importantly, while random death can occur anywhere within the tumor (see blue population in Figs. 1C and 1E), the death by drug exposure is assumed to only penetrate to the cells on the periphery (see black population in Fig. 1E) as in our previous model [44]. The periphery of the tumor is dynamic as treatment is applied and withdrawn and random death within the tumor further complicates this delineation, so the tumor periphery is determined using the occupation of neighboring cells and grid spaces. This is explained in detail in the Supplemental Methods. In prostate adaptive cancer therapy, abiraterone acetate (an antiandrogenic CYP17A inhibitor) is administered adaptively in combination with continuous administration of leuprolide (androgen deprivation therapy). However, to extend the modeling framework to consider applications of adaptive therapy beyond prostate cancer, it is important to consider a range of treatment mechanisms and each mechanism’s effect on adaptive dynamics. In particular, we explore two theoretical drug mechanisms: 1) cell-cycle independent, which occurs at a specific rate at every time step, and 2) cell-cycle dependent, which acts only upon proliferating cells during cell division (see Fig. 1F). The probability of cell death per time step for the cell-cycle *independent* drug is:

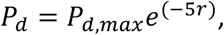

where P_d,max_ is the maximum probability of cell death per day, and *r* is the level of resistance. Here, we assume that resistant cells have a fitness cost, in that their rate of division is slower than that of a fully sensitive cell. Specifically, the level of resistance follows a linear tradeoff with the proliferation rate (see range in Table S1) and is calculated as *r* = 1 - *p*, where *p* is normalized from 0-1. The probability of cell death per division by the cell-cycle *dependent* drug is:

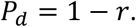

In both cases, dead cells are removed after 18 hours [44].

#### Simulation Environment

The model was programmed in the Java language (ver. 11.0.2), using the Abstract Window Toolkit for visualization. The platform for the program was an Intel Core i7-8559U processor as well as the high-performance computing cluster at H. Lee Moffitt Cancer Center and Research Institute (Tampa, FL). The source code is available on Github at: https://github.com/jillagal/multiMets.

## Results

The aim of this paper is to investigate whether the dynamics during the initial cycle of adaptive therapy can be used to learn about the characteristics of the underlying disease. To begin with, we characterize the cycling dynamics in adaptive prostate cancer therapy patients.

### Regrowth times are typically longer than treatment response times in the first drug cycle

We analyze the mCRPC trial patients to observe the variability in the dynamics of the first treatment cycle, which is the period of first applying therapy, removing it upon 50% PSA reduction, and regrowth until initial PSA levels are reached. We find that there is a range of cycle times (Fig. 2A), with differences in response times (when the treatment is on), and in the regrowth times (when the treatment is off). Further, we can visualize this initial cycle in “cycle space”, by plotting the response time on the *x*-axis and the regrowth time on the *y*-axis for each patient (Fig 2B). While there is variation between patients, we note that on average, the time that the drug is applied is shorter than the time that it takes the PSA to return to the original value after the drug is withdrawn (i.e. the points mostly lie above the diagonal). To illustrate this for individual patients, we show the PSA levels over time for three patients distributed across this cycle space (Fig. 2C).

**Figure 2.**
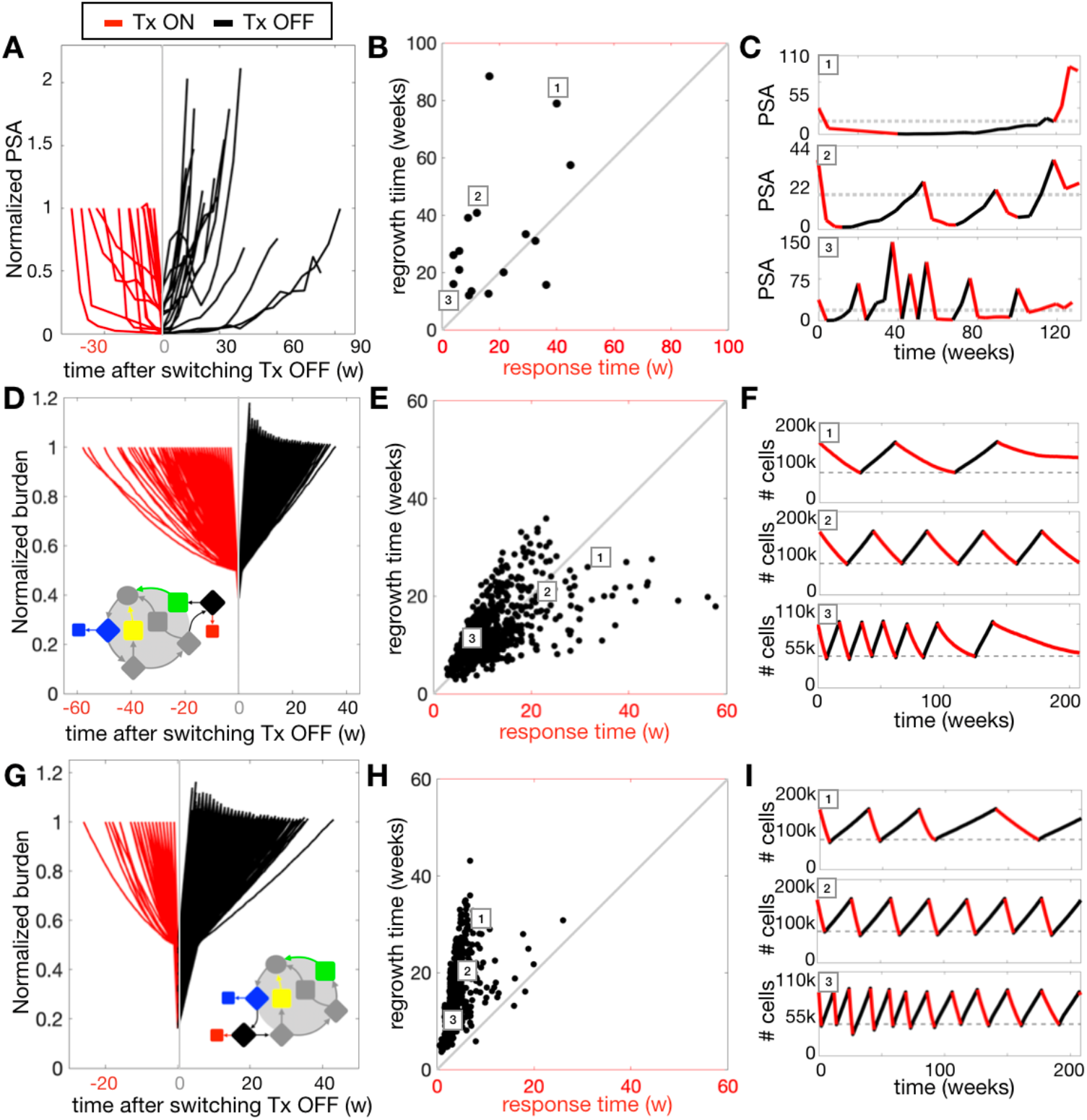
Timing variability of adaptive therapy cycles in patient data and the mathematical model. A-C) Patient data (n=16) from the mCRPC adaptive therapy trial, D-F) model results using a cell cycle-dependent drug, and G-I) model results using a cell cycle-independent drug. For each row of panels, the left column shows a timeline of the first on/off treatment cycle for all patients/simulations in the cohort. The burden is normalized at the initial value when treatment starts, and the timeline is shifted so that the zero-time point (gray line) is at the switch point from treatment on to treatment off. On-treatment cycle times (Tx ON) in red are flipped about the y-axis for visualization purposes. The middle column shows the “cycle space”, where there is a point for each patient/simulation showing the first treatment cycle with the treatment response time on the *x*-axis and the regrowth time on the *y*-axis. The numbered boxes in the cycle space point to specific patients/simulations within the space, which are displayed in the right column as a timeline over the full course of treatment. The three specific example simulations in F and I share the same parameter sets: 1) [N_Dx_, m, r_0_, d_r_, h_i_, h_a_]=[150000, 1, 0.4, 0.06, 0.5, 0], 2) [N_Dx_, m, r_0_, d_r_, h_i_, h_a_]=[150000, 1, 0.2, 0.06, 1, 0.5], and 3) [N_Dx_, m, r_0_, d_r_, h_i_, h_a_]=[80000, 10, 0.4, 0.033, 1, 0.5].

### A cell cycle-independent drug is needed to account for short response times and longer regrowth times

To explore the impact of different parameters on the treatment dynamics, we create a cohort of 1000 virtual patients by randomly varying the six key model parameters shown in Table 1 and running each simulation through the growth/seeding period. The final states of each of these 1000 sets of metastases become the initial conditions for two parallel treatment arms: a cell cycle-dependent drug and a cell cycle-independent drug. For both drugs, we find variation across the simulations (Figs. 2D-I). Under the cell cycle-dependent drug treatment (Fig. 2D-F), the response and regrowth times are nearly equal (10.2±6.9 and 11.1±6.2 weeks, respectively, for mean and standard deviation). This can be explained by the fact that death can only occur as fast as division when the drug is on, while regrowth occurs at that same rate of division. There are also some simulations that suggest the possibility of slow response rates followed by fast regrowth. This variability is due to stochastic effects, spatial effects, and parameter effects that are explored later. In contrast, the cell cycle-independent drug kill (Fig. 2G-I) on average has shorter response times and longer regrowth times (3.5±2.1 and 14.0±5.9 weeks, respectively).

**Table 1.** Ranges of parameters used in the mathematical model.

| Parameter | Description | Range |
| --- | --- | --- |
| $N_{Dx}$ | Total number of cells at diagnosis | $10\text{-}150 \times 10^3$ |
| $m$ | Number of metastases seeded | 1-10 |
| $r_0$ | Initial mean drug resistance | 0-0.4 |
| $d_r$ | Random death rate | $0\text{-}6 \times 10^{-2}/\text{day}$ |
| $h_i$ | Intertumor heterogeneity | 0-1 |
| $h_a$ | Intratumor heterogeneity | 0-1 |

A visual comparison of the cycle space for the cell cycle-dependent drug (Fig. 2E) and cell cycle-independent drug (Fig. 2H) implies that the cell-independent drug is more representative of the skewed relationship between response and regrowth times seen in the patient dynamics (Fig. 2B). We hypothesize that this is because the cell cycle-independent drug model is a closer representation of the drug action of abiraterone, which acts to stop production of testosterone, a necessary resource for most prostate cancer cells. Example simulations are shown (Figs. 2F and 2I) and demonstrate the representative differences over different cycle lengths. Given its superior ability to describe the patient data, we focus on the cell cycle-independent drug application in the remainder of this paper.

### Smaller metastases have shorter drug cycles

To gain an understanding of how the metastatic distribution affects treatment cycles, we look at how cycles are affected by the total tumor burden and the number of metastases present at the start of treatment. We create a new virtual patient cohort by randomly sampling 300 values of burden and initial number of metastases (from the ranges in Table 1), while keeping the other parameters fixed. First, we assume that drug resistance is 0, cell turnover is 0, and the tumors are completely homogeneous. We observe that the shortest response times (Fig. 3A) happen when there are many metastases and a lower total burden, while the longest times occur when there is a large burden and few metastases. The regrowth times follow a similar pattern but with slightly longer times (Fig. 3B). We isolate several examples in this space (labeled 1-4). Examples 1 and 2 both have 2 metastases and either a high or low burden, respectively, while examples 3 and 4 both have 9 metastases with either a high or low burden, respectively (Fig. 3C, left). Example 1 has the longest treatment cycles, examples 2 and 3 have similar cycles, and example 4 has the shortest treatment cycles (Fig. 3C, right). The reason for this difference lies in the spatial constraints on tumor growth. A large total burden split into a few metastases results in a few large lesions, while the same burden split into many metastases results in many smaller lesions. With a larger burden per metastasis, the perimeter-to-area ratio decreases, slowing the response and regrowth times, since the larger metastases must lose and regrow a larger number of cells to reach the 50% and 100% treatment switching thresholds.

**Figure 3.**
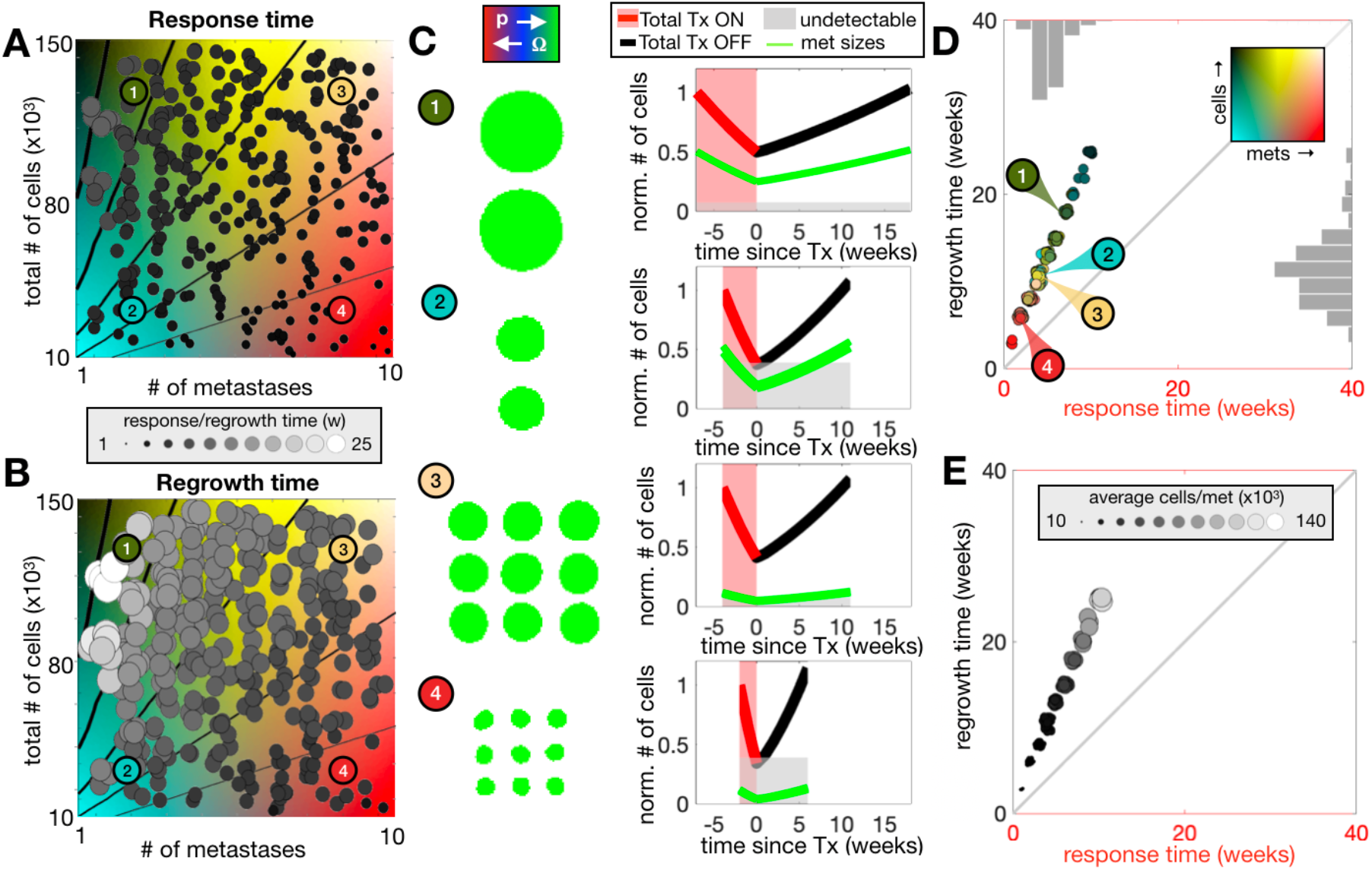
The effect of total tumor burden and number of metastases on adaptive therapy cycling. An array of 300 random parameter combinations are simulated and shown colored in grayscale according to A) drug response time and B) regrowth time, which both use the same array of values and have added noise for easier visualization. Isolines are shown at 5k, 10k, 20k, 40k, and 80k cells per metastasis with increasing line weights. C) Example simulations with parameter combinations 1 through 4 (as labeled in A and B). The green tumors represent the multiple lesions pre-treatment, while the plots show the dynamics of the lesions during the first cycle of adaptive therapy. The total burden is in red (drug on) and black (drug off) and the individual metastases are plotted in green. The light gray rectangle shows our chosen threshold of metastatic invisibility at 10k cells. D) The same simulations from panels A and B are projected into cycle space and colored according to their combination of total burden and number of metastases using the background colormap. The histograms show the frequency distribution of each axis separately. E) We plot the same points from D colored according to the average number of cells per metastasis for each simulation. [N_Dx_, m, r_0_, d_r_, h_i_, h_a_]=[10000-150000, 1-10, 0, 0, 0, 0].

As such, the cycle times do not depend on the number of metastases or total burden, but the average size of the individual metastases (see contour lines of constant metastases size in 3A-B). To illustrate this, we plot the cohort in cycle space and color each parameter combination by the number of metastases and the total tumor burden. This shows that the full range of metastatic number and size can be projected along a single axis in cycle space, where certain parameter combinations give the same cycling dynamics (Fig. 3D). When we instead color by the average number of cells per metastasis (Fig. 3E) there is a clear pattern showing that the average size of the metastases determines the position in cycle-space. To sum up, larger metastases have longer cycles, and smaller metastases have shorter cycles.

### Drug resistance and turnover affect treatment cycles

Next, we look at how treatment cycles are affected by death parameters in a new cohort of virtual patients with a range of drug resistance and cell turnover rates (see Fig. 4, Table 1). In this cohort, two homogenous metastases are grown to a combined burden of 80,000 cells prior to treatment. The results show that drug response time is shortest when resistance is low and turnover is high, while the response time is longest when the opposite is true (Fig. 4A). Generally, regrowth times are quicker when resistance and turnover are low (Fig. 4B), and slower when the opposite is true. The examples (1-4) examine the mechanisms that cause these trends (Fig. 4C). The effects observed with resistance are straightforward: highly resistant examples (3 and 4, blue lines) will respond slower to the drug (longer drug application periods), and also take longer to regrow (longer drug-free periods) because of the assumed resistance-proliferation tradeoff. The effect of turnover is slightly more complicated. Examples 1 and 3 have an increased number of proliferating cells (green cells), due to increased open space from high random death. This increases drug penetration and shortens the drug application times. During the drug-free regrowth period, random cell death slows the net growth rate, increasing regrowth times. These effects contribute to how different combinations of these parameters affect a tumor’s cycle times during adaptive therapy (Fig. 4D). In the previous section, we noted that the overall cycle time is mostly determined by the tumor size. Given that these tumors are essentially the same size when treatment begins, we see that the combination of drug resistance and turnover contributes to further variation around that position within the cycle-space, which is skewed slightly toward shorter response times. Additionally, the combination of effects on growth rate by these parameters leads to different net growth rates, which can be observed by coloring the same points from Fig. 4D in terms of tumor age at diagnosis (Fig. 4E). The older metastases take longer to reach the same size and have longer treatment cycles while the younger metastases have shorter treatment cycles.

**Figure 4.**
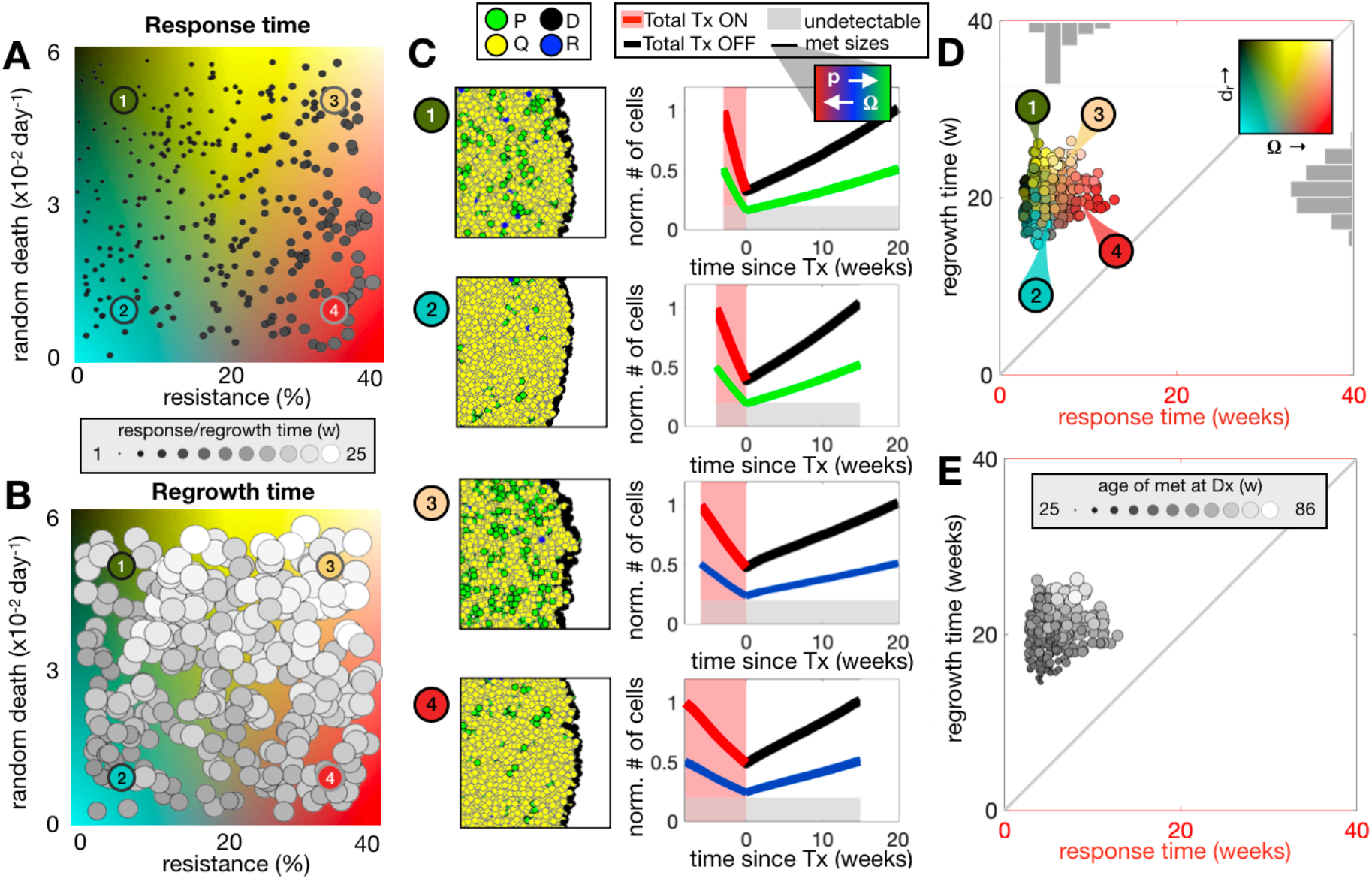
The effect of the level of pre-existing drug resistance and turnover on adaptive therapy cycle dynamics. An array of 300 random parameter combinations are simulated and shown colored in grayscale according to A) drug response time and B) regrowth time. C) Example simulations with parameter combinations labeled in A and B are shown colored according to their phenotypic state (P=proliferating, Q=quiescent, D=druggable, R=random death) and over the first cycle of adaptive therapy. The dynamics show the total burden in red (drug on) and black (drug off) and the individual metastases in green. The light gray rectangle shows our chosen threshold of metastatic invisibility at 10k cells. D) The same virtual cohort in A) and B) are plotted in cycle space and colored according to their combination of sensitivity and random death rate parameters using the colormap from A and B. E) The same points from D are colored according to the age of the metastases prior to treatment for each simulation. [N_Dx_, m, r_0_, d_r_, h_i_, h_a_]=[80000, 2, 0-0.4, 0-6 ×10^−2^, 0, 0].

### Seeding asynchronicity and intertumor heterogeneity affect tumor size distributions

We previously assumed that all metastases were seeded simultaneously and that metastases were homogeneous. This means that despite some stochasticity, all tumors within the system of metastases were roughly the same size. Here we investigate the impact of having differently sized metastases, particularly those below the threshold of detectability. Metastases may be undetectable due to a slow growth rate or late seeding; therefore, we compare two possible scenarios for metastatic seeding: 1) homogeneous sequential seeding and 2) heterogeneous simultaneous seeding. For 1), we seed the first tumor at time 0 and the remaining tumors are seeded at a constant rate throughout the growth period prior to treatment. All tumors are the same phenotype, delayed in time. For 2), we seed all tumors simultaneously at time 0. In this case, each lesion is different, composed of a single phenotype of cells drawn from a range of proliferation rates (and therefore a range of treatment sensitivities). The amount of this “intertumor heterogeneity” can be set between 0 (all lesions are the same) and 1 (proliferation rates are selected from the full range of the values permitted for proliferation from Table S1). All other parameters are fixed (Table 1).

To compare with the previous results, we show simulations using homogenous tumors seeded simultaneously across a range of total burden and number of metastases (Figure 5, left column). We plot the number of visible metastases within this parameter space. As above, because all metastases are seeded simultaneously and are homogenous, they are similar in size at time of therapy. Therefore, we find that generally either all of the metastases are visible, or all of them are below the threshold and therefore not visible (Fig. 5A). Similarly, for sequential seeding we find that when there are only a few metastases with a large burden, they are all visible, and when there are many metastases making up a small burden, they are all undetectable (Fig. 5B). However, with a larger burden spread amongst more metastases of various sizes, some of the late-seeded, smaller metastases may not be observed. A similar pattern occurs with intertumor heterogeneity (Fig. 5C). We find here that the high burden, high number of metastases portion of the plot has expanded substantially so that only the single metastases or several very large metastases are completely accounted for, whilst most simulations have at least some undetectable metastases.

**Figure 5.**
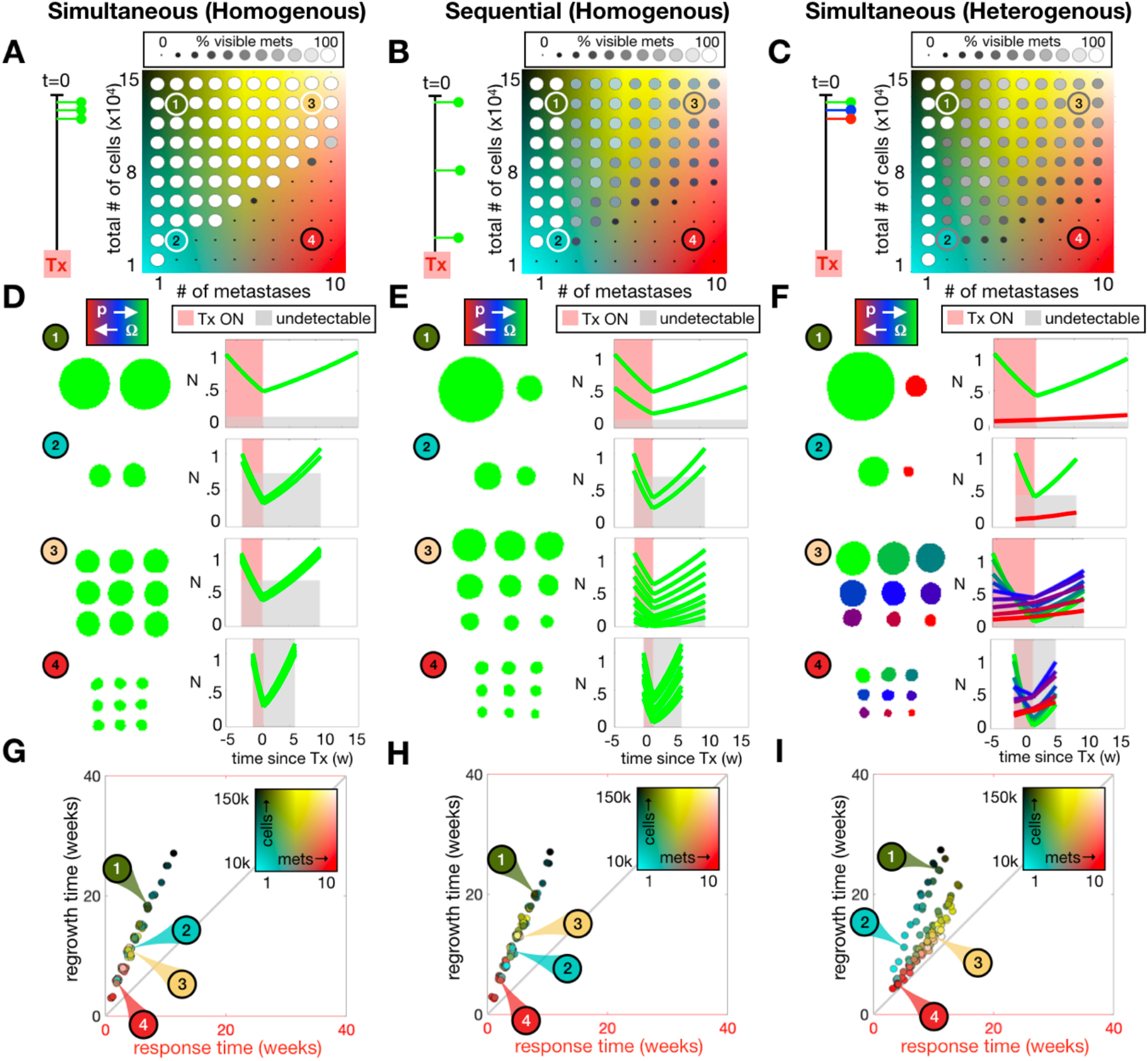
Comparing effects of seeding timing and intertumor heterogeneity on metastasis visibility and cycle times. Metastases are either seeded A) simultaneously, B) sequentially, or C) with heterogeneous phenotypes simultaneously. For each case, an array of values for total tumor burden and number of metastases are plotted colored according to the % of visible metastases. D-F) Spatial distributions of several example metastases are shown prior to treatment, and temporal dynamics of individual metastasis sizes are shown over one cycle of adaptive therapy. G-I) The same simulations from the arrays in A-C are plotted in cycle space using the same background colormap that designates the combination of number of metastases and total number of cells. *[N*_*Dx*_, *m, r*_*0*_, *d*_*r*_, *h*_*i*_, *h*_*a*_*]=[10000-150000, 1-10, 0, 0, 0, 0]*. Threshold of visibility is set at 10,000 cells.

Intriguingly, Fig 5D-F shows that while both homogeneous sequential seeding and heterogeneous simultaneous seeding results in a distribution of sizes of metastases, some which are below the threshold of detectability, the changes of the tumor sizes over a cycle of adaptive therapy can be informative. Several examples in this parameter space are displayed spatially prior to treatment and the normalized sizes of metastases are shown over one cycle of adaptive therapy (Fig. 5D-F). With simultaneous, homogenous seeding all tumors are nearly the same size prior to the treatment cycle as after the treatment cycle (Fig. 5D). In the sequentially seeded tumors (Fig. 5E), whilst there is size variation amongst the lesions prior to treatment, they also return to a similar size distribution after treatment. There are also different sized tumors within the simultaneous heterogeneous metastases (Fig. 5F). The most significant difference between sequential seeding and heterogeneous seeding is that the most resistant tumors will not respond to treatment. Therefore, while the larger more sensitive tumors dominate the dynamics at the start of the treatment cycles, smaller more resistant tumors will grow unimpeded during treatment and may contribute more to the burden over the course of the cycle (e.g., Example 4 in Fig. 5F).

Furthermore, we find that the size and phenotype heterogeneity amongst the metastases affects the position in cycle-space (Fig. 5G-I). Ultimately, when the cycles for the sequentially seeded tumors are plotted in cycle-space (Fig. 5H), we see little difference from the baseline, but some of the cycle times have shifted (compared to Fig. 5G). The metastases in Examples 2 and 4 (Fig. 5H?) are grown to a small burden and have very little difference between the oldest and the youngest metastasis, compared to those in Fig. 5G. However, for Examples 1 and 3 that are grown to a larger total burden, the earlier seeding results in a more significant size difference of the largest tumors, which is enough to elongate the cycle times for that set of metastases. We therefore conclude here that it is not actually the average size of the metastasis that affects the cycle time but rather the largest tumor dominates the dynamics. For the heterogeneous metastases (Fig. 5I), the largest tumor still dominates the cycle times in much the same way if there is a large difference in size of the metastases (Examples 1 and 2). However, when more resistant cells make up a significant portion of the total burden (Examples 3 and 4), cycle times are elongated for both response and regrowth, due to slow-growing drug resistant cells.

### Tumor heterogeneity affects treatment response

In the previous sections, we showed that drug resistance affects cycle times. Further, heterogeneity amongst the metastases affects cycle times. Here, we examine how different types of heterogeneity affect the outcome and if a continuous dose would be a better option for each virtual patient in the cohort. For this question, we tested an adaptive therapy (AT) schedule against continuous treatment (CT) to consider which situations would respond better with the aim of tumor control as opposed to tumor elimination. We find that most of the simulations result in an eventual recurrence, but in 35% of the cohort, continuous treatment leads to tumor elimination. The tumors that are eliminated, on average, have both low intertumor and intratumor heterogeneity compared to those that recur (Fig. 6A). Of those that do recur, we compare their time to recurrence (TTR) on each treatment schedule with respect to measured heterogeneity prior to treatment (Fig. 6B-C). We plot the TTR for AT on the *x-*axis and for CT on the y-axis, so that for each pair of treatments a point on the line means that there is no difference in TTR for CT and AT, while a point above the line favors CT and a point below the line favors AT. The majority (82%) of simulations lie below the line, favoring an AT schedule. With some exceptions, the cases with the most intertumor heterogeneity respond better to continuous treatment (Fig. 6B) while the cases with more intratumor heterogeneity respond better to adaptive therapy (Fig. 6C). To understand why, we consider two extreme scenarios that are initialized with maximum intertumor heterogeneity with no intratumor heterogeneity and maximum intratumor heterogeneity with no intertumor heterogeneity. When the tumors with intertumor heterogeneity (Fig. 6D) are treated continuously, the total burden and the number of metastases decreases, but ultimately, the total burden starts to increase again as the most sensitive tumors are killed off and the more resistant tumors take over the dynamics. If instead adaptive therapy is applied, partially responsive tumors continue to grow when they could have been killed by ongoing treatment. In this scenario, the benefit of having sensitive cells to suppress the growth of resistant cells is not applicable because they are not competing for the same space, being in different lesions. However, with intratumor heterogeneity (Fig. 6E), a mixture of phenotypes within each metastasis do compete for the same space, so that adaptive therapy can delay treatment failure. In contrast, when continuous treatment is applied to this patient, there is competitive release of resistant phenotypes, which results in earlier recurrence. Ultimately, these results show that a CT or AT treatment strategy may be preferred to extend TTR for patients with high inter- or intra-tumor heterogeneity, respectively. However, for patients with low heterogeneity, the challenge is that cure may be possible using CT, while the best AT can do is continuous control. All in all, AT offers less risk: if cure fails, then AT is preferable to extend TTR, and the time gained by using AT when it is optimal far exceeds the time lost when using AT when CT is optimal.

**Figure 6.**
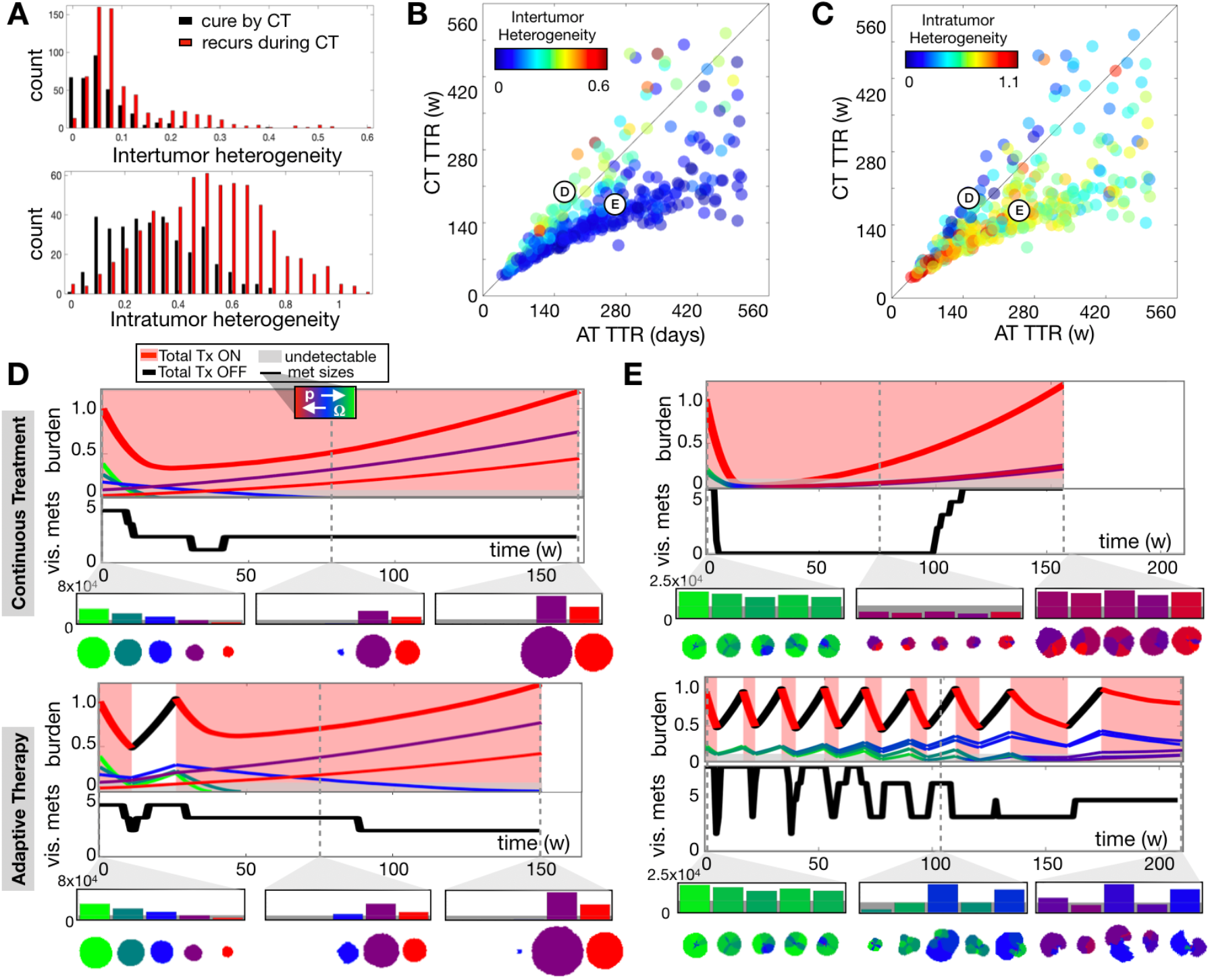
Comparing responses to continuous treatment (CT) vs adaptive therapy (AT) in the model cohort (n=1000). A) Histograms of simulations with different values of intertumor heterogeneity (top) and intratumor heterogeneity (bottom) that respond in a complete (cure) or incomplete (recurrence) response to continuous treatment. B-C) Time to recurrence (TTR) under CT and AT for each simulation, colored according to intertumor heterogeneity (B) and intratumor heterogeneity (C). Gray line indicates when there is no difference in the time to recurrence between the two treatments. D-E) Examples of metastases with intertumor heterogeneity (D) and intratumor heterogeneity (E) comparing CT (top) and AT (bottom). For each show systemic burden and individual metastases over time (top panel), the number of visible metastases (>10k cells) over time (middle panel), and a histogram of the metastases size distribution with each colored according to its average phenotype (lower panel). The corresponding spatial distribution of each metastasis is shown below each bar.

### Changes in visible metastases indicate intertumor variability

Having characterized how different tumor parameters affect the adaptive therapy cycling dynamics, we apply our novel understanding to the patients in the clinical trial [9]. In the above results, we have examined how tumor parameters in isolation or in pairwise combinations alter the adaptive therapy cycling dynamics. However, metastatic disease is a complex system and patients are likely to differ in multiple parameters at once. In the final part of this paper, we therefore aim to relate parameter combinations back to the patients in the trial using the Gleason score [45] of the primary lesion, which correlates with a number of clinically important features such as outcome. In this cohort, we observe three distinct categories of response, which we will refer to as low Gleason (6-7), mid Gleason (8), and high Gleason (9-10). Over the first cycle of adaptive therapy, patients with low Gleason scores have either the same or fewer metastases at the end of the cycle than at the start of the cycle (Fig. 7A). In contrast, all patients with mid Gleason Scores have no change in the number of metastases over the first cycle, and patients with high Gleason scores have either the same or an increase in the number of metastases. Further, the mid Gleason patients are seen to have a higher average number of cycles over the full treatment course, while low and high Gleason patients have only a few (Fig. 7B). Finally, the patients with the mid Gleason scores tend to have the fastest cycle times (Fig. 7C).

**Figure 7.**
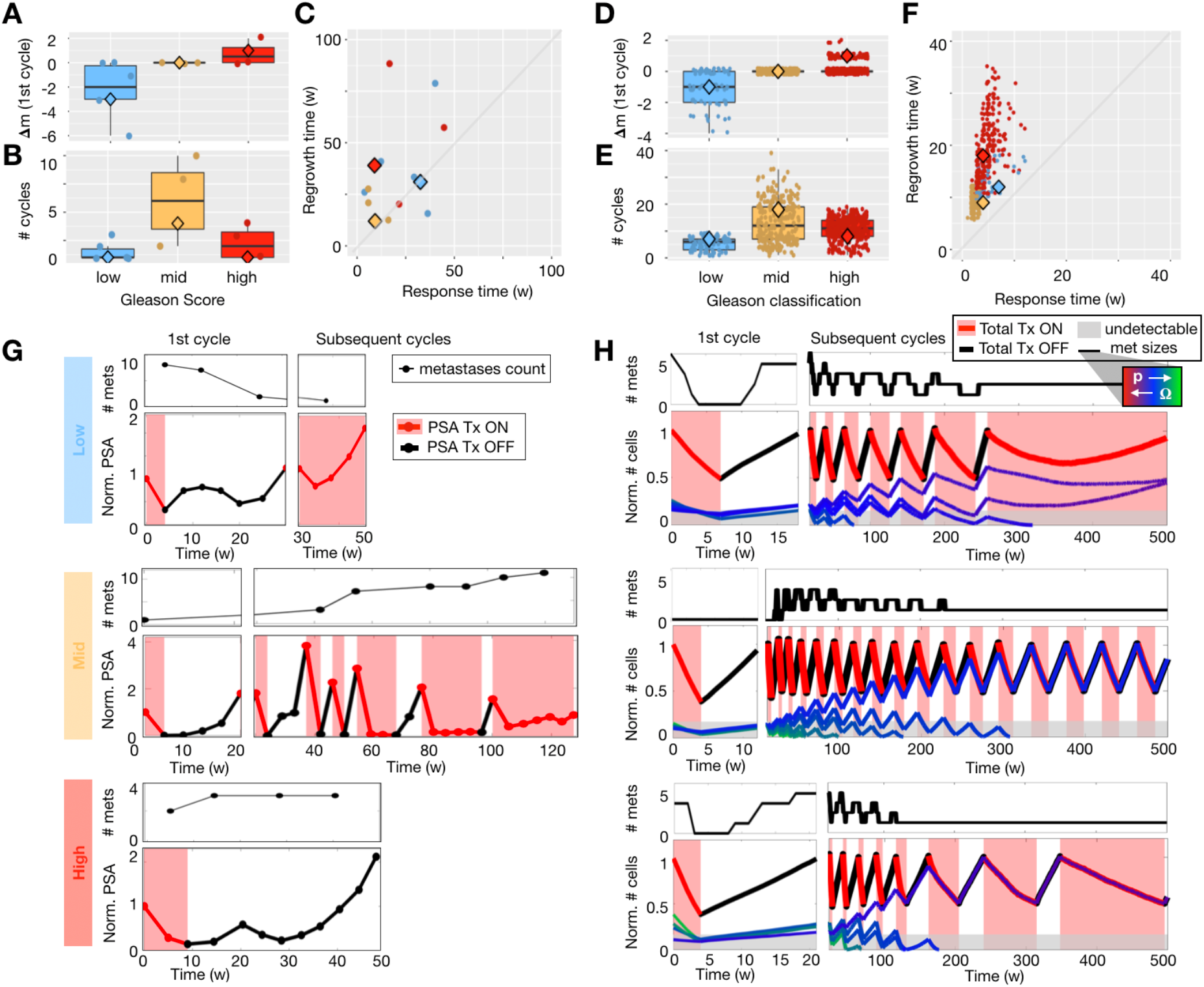
Comparing patient metrics (n=14) grouped by Gleason Score (low=6-7, mid=8, and high=9-10) to simulation data. Patient data is shown in A-C & G, and model data is shown in D-F & H. A) Relationship between Gleason Score and the change in the number of metastases (A), the total number of treatment cycles (B), and the patient’s location in cycle space (C). Note that 2 patients were left out of this analysis due to one missing Gleason Score and one with insufficient data on the number of metastases. D-F) Model data from Fig 2E & F classified into a Gleason category based on the change in the number of metastases, the number of cycles, and the timing of their cycles (algorithm explained in the Supplemental Methods; patients that were unable to be classified are not shown). Large diamonds represent the patients and simulations that best fit the average metrics of each Gleason group. G) Treatment dynamics of patients best representing the average of each Gleason group, and H) simulations predicted by the model for the best fits. These correlate with the large diamonds in A-F.

Next, we wanted to assign a Gleason score to the virtual patients in the cohort of 1000 simulations used in Fig. 2C-F (where all parameters were varied). While we have no equivalent to the Gleason score in the computational model, we can match simulations with a putative Gleason score based on comparative metrics to the patient data from Fig. 7A-C, which include the change in number of metastases over the first cycle, the total number of cycles over the course of therapy, and the response and regrowth time of the first cycle (see Supplemental Methods for details). The distributions and individual values for each metric in the simulated cohort classified by Gleason group are shown in Fig. 7D-F. We were able to classify 68% of the simulations with 8.2%, 26.5%, and 33.3% in the low, mid, and high Gleason groups, respectively. The rest of the data did not fit into any group and is not shown. The patient treatment trajectories for best fit to the average metrics are shown in Fig. 7G. We observe that the patient with a low Gleason score has a decrease in the number of metastases over the first cycle and fails shortly after that. The patient with a mid Gleason score has no change in the number of metastases during the first cycle and continues with six more relatively quick cycles. The patient with the high Gleason score has an increase in metastases during the first cycle and was taken off the study at that point. The model-predicted treatment trajectories for the best fitting simulation for each Gleason group are shown in Fig. 7H. The simulations with both high and low Gleason classifications had longer cycling times during the first and subsequent cycles, while the simulation with the mid Gleason classification had faster cycles with no change in the number of metastases over the first cycle.

While these patients and simulations represent the best fit to the average of the data, it is important to note that there is variation amongst key qualitative features within the group like outcome. In other words, many cycles does not mean longer time to recurrence, but the model suggests that it may mean smaller lesions. We examined what other metastatic features our model fits could tell us about the virtual cohort (Fig. S2). We find a noticeable difference in heterogeneity between the Gleason groups. We calculate both intertumor and intratumor heterogeneity just prior to treatment as the standard deviation of the mean phenotype for each lesion and the mean of the standard deviation of phenotypes for each lesion, respectively. The mid Gleason groups are found to have the lowest intertumor heterogeneity and the highest intratumor heterogeneity. This result illustrates how integrating the data from adaptive therapy cycling with mathematical modelling may help us to better characterize a patient’s disease.

## Discussion

Here we introduced a multiscale model to map tumor dynamics onto cycle space, showing that patient-specific response and regrowth times can imply disease characteristics like metastatic distribution, resistance, and optimal therapy strategies. Characterization of metastases is inherently challenging due to dispersion of metastatic seeds to many sites and the sparsity of data collected beyond just presence or absence of metastases. However, vigilant observation during a single adaptive therapy cycle could provide new knowledge. With further analysis and validation of the metrics introduced here, they may add supporting evidence to existing biomarkers and aid in personalizing treatment for patients with metastatic cancer.

During a single cycle of adaptive therapy consisting of a drug application period and a regrowth period, we can delineate multiple parameters of the metastatic system. We found that with the same initial burden, more metastases resulted in quicker adaptive therapy cycle times. However, it is not the number of metastases that matters, but the largest lesions that dominate the dynamics. Larger metastases have longer cycles, while smaller metastases have quicker cycles, similar to findings where more spread out tumor seeds had faster cycles than larger patches of cells in [18]. We also found that more drug resistance slows the treatment cycles, as expected, and a faster cell turnover speeds up drug response time by increasing the surface area to aid in drug penetration but slows the regrowth time by reducing the net proliferation rate. The latter result was also found in a non-spatial model [19], where turnover increased total cycle times, which are generally dominated by long regrowth periods and shorter response times.

It is also important to recognize the limitations of imaging for characterizing metastases. A single snapshot in time may not capture smaller, later-seeded lesions. However, variation between metastases with regards to size and phenotype may be captured in the cycle dynamics. We found that the cycle dynamics are dominated by the largest tumor, but if enough resistant tumors exist in a large enough proportion, the cycle times will increase. Further, the change in the number of metastases over the cycle rather than just a single time point may yield more information. We found that change in metastases over the course of a cycle may indicate that there are differences amongst the metastases, which could be associated with the Gleason score of the primary tumor. Both lower Gleason scores (6-7) and higher Gleason scores (9-10) show more cases where the number of metastases decrease and increase, respectively, during the first cycle of adaptive therapy. These two groups also tend to have very few total cycles that are longer when compared to mid Gleason scores that on average have more cycles that are shorter duration. Classification of the model results into Gleason groups in this model revealed it may be due to differences in phenotypic heterogeneity, with respect to proliferation rate and treatment resistance. More intertumor heterogeneity was found in low and high Gleason grades, which ultimately causes treatment failure due to failure from specific lesions, and more intratumor heterogeneity was found in mid Gleason grades, which allows for local competition amongst phenotypes within a single lesion that is necessary for the success of adaptive therapy. It has otherwise been shown that mid Gleason grades are unique in that the cells may be in a transitional state [46], which may aid in treatment response. However, the increased response of mid Gleason grades to adaptive therapy does necessarily correlate with outcome. While the standard-of-care continuous treatment does show that higher Gleason grades have a shorter time to progression, with adaptive therapy the time to progression somewhat evens out over the Gleason grades (Fig. S3). In this analysis, we have used large scale clinical features from patients with known Gleason grades to stratify systems of computational lesions with known cell-scale phenotypic characteristics into corresponding Gleason grades. However, it should be noted that the number of patients in the clinical trial is small, so these results should be further explored for validation.

Here we have considered a simplified version of a system of metastases to get a basic understanding of individual and systemic dynamics during one cycle of adaptive therapy. The aim was to capture generic elements of metastases during treatment while comparing to the mCRPC trial for guidance. We have neglected some important elements such as the effect of early or late seeding [47]– [49], metastases seeding metastases [50], differences in metastasis cluster size [51], [52], dormancy [53], [54], diversity of metastatic potential [51], [55], [56], drug-induced resistance at the metastatic site [57], [58], and stromal interactions in metastatic niches [46]. We also modeled a tradeoff between drug resistance and proliferation, which may not always apply. In addition, the time and space scales were both small compared to those in the mCRPC trial. The size scale was limited by the domain size, which was kept small for computational efficiency, and the timescale was reduced in turn. Therefore, cycling times were smaller, and we needed to use 1-week intervals to check burden and make treatment decisions rather than the 4-week intervals in the clinical trial. These scale issues could be diminished by using a different, coarser model system, but would likely come at the expense of not having cell-scale spatial interactions that may be important mechanisms affecting the outcome, due to heterogeneity and competition within a lesion.

Critically, combining available metrics on different scales of resolution may aid in a better characterization of disease at this late stage when metastases may be invisible and challenging to quantify, as each modality has limited scope in isolation. Blood biomarkers are not perfect, yet they remain a cheap viewpoint of a systemic measure of tumor burden over time. Imaging also has resolution limits, but changes over the cycle may help inform treatment choice. Ultimately, there are only two reasons why continuous, maximum dose treatment succeeds: the metastases do not have pre-existing heterogeneity, or the harsh treatment prevents the resistant phenotypes from taking over. However, when treatment fails, it can do so in different ways. Most simulations that resulted in recurrence favored an AT schedule. However, some cases with the most intertumor heterogeneity responded better to continuous treatment. Perhaps a blood biomarker signature could be developed to quantify tumor heterogeneity that when combined with these results could make more informed clinical decisions. Currently, cancer staging provides only a binary view of metastatic presence or absence. Variations on metastatic staging metrics only give a coarse-grained resolution of an already puzzling view of late-stage disease, so identifying additional metrics based on tumor dynamics could be a step forward. While this work is only a first step, it shows that careful monitoring of biomarkers and visible metastases during one cycle of adaptive therapy may be useful in identifying important features of the metastatic structure to provide further support for treatment decision making.

## Supplement

### Supplemental Methods

#### Calculate druggability of a cell

Instead of modelling the dynamics of the drug explicitly, such as diffusion, uptake, decay, and perfusion into the tumor, we made the simplifying assumption that the drug will only reach the outer periphery of the tumor. In previous versions of this model [37], this was achieved by only allowing the proliferating cells access to the drug, which were always on the boundary due to quiescence when there was no space to divide. However, adding cell turnover means that cells can die in the middle of the tumor, which opens up space for cells to reenter the proliferation state and therefore be exposed to the drug. Here, we apply the following algorithm for ensuring that only the cells on the periphery of the tumor will be subjected to drug exposure.

At the start of each cell loop, cells are assigned a position in a coarse grid (18×18 pixel grid over a 2700×2700 pixel domain) to determine an approximate neighborhood, which is used to narrow the range of possible neighbor interactions and speed up computation. This same grid structure is used to determine if a cell is on the periphery of the tumor. If the cell is on the periphery, it will be druggable. The peripheral cells are those cells that are on the edge of the mass of cells, which we define explicitly by either 1) there is more than 1 neighboring grid point that has less than 2 cells, or 2) there is only 1 grid point with fewer than 2 cells but those cells are at least a half of a grid length away from the cell in question. Further, the cell must be in the proliferating state to be affected by the drug.

#### Fixed parameters

**Table S1.** Other parameters that do not vary.

| Description | Range | Reference |
| --- | --- | --- |
| proliferation rates | 0.12-0.48 d<sup>-1</sup> | estimated |
| cell diameter | 20 μm | estimated |
| probability of cell death by cell cycle-independent drug | 0.6 d<sup>-1</sup> | estimated |
| Initial number of cells in metastasis seed | 30 | set |

##### Gleason Classification

The simulation data are classified into a Gleason group (high, mid, and low) based on how several metrics compare to the those from the patient data (from Fig. 6A). First, the virtual patient (simulation) data are grouped into a possible Gleason group based on the change in the number of metastases over the first cycle of adaptive therapy. If there is a decrease in metastases, the virtual patient may only be in the low Gleason group. If there is no change in the number of metastases over the first cycle, it may be classified into any group, and if there is an increase in metastases, it may only be in the high Gleason group. Further, we check the probability that each virtual patient may fit in either group based on the distributions of the other 3 metrics (number of total cycles, drug response time, and regrowth time). Because the patient’s tumors sizes are on a much larger scale than those simulated, the time scales will be shorter. This will change the scale of all 3 metrics that we are comparing. Therefore, we do not compare absolute values, but rather we normalize the values in both the patient data set and the simulated data set by their maximum to compare. We assume that the distribution of each metric follows a normal distribution so that we can compute the probability density that a value xi from the simulated data lies within the distribution of the metric *i* as:

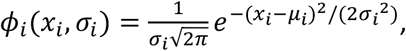

where µ_I_ and α_I_ are the mean and standard deviation of the patient distribution. Then for each virtual patient, we calculate the probability that it belongs to a Gleason category as the mean of probabilities over the 3 metrics. If the probability is below 70%, it may exist in that group. If it can exist in multiple groups, it will be classified into the group that has the highest probability. If it does not meet any of these criteria, it will be deemed as not belonging to any group and will be removed from the analysis.

## Supplemental Results

**Figure S1.**
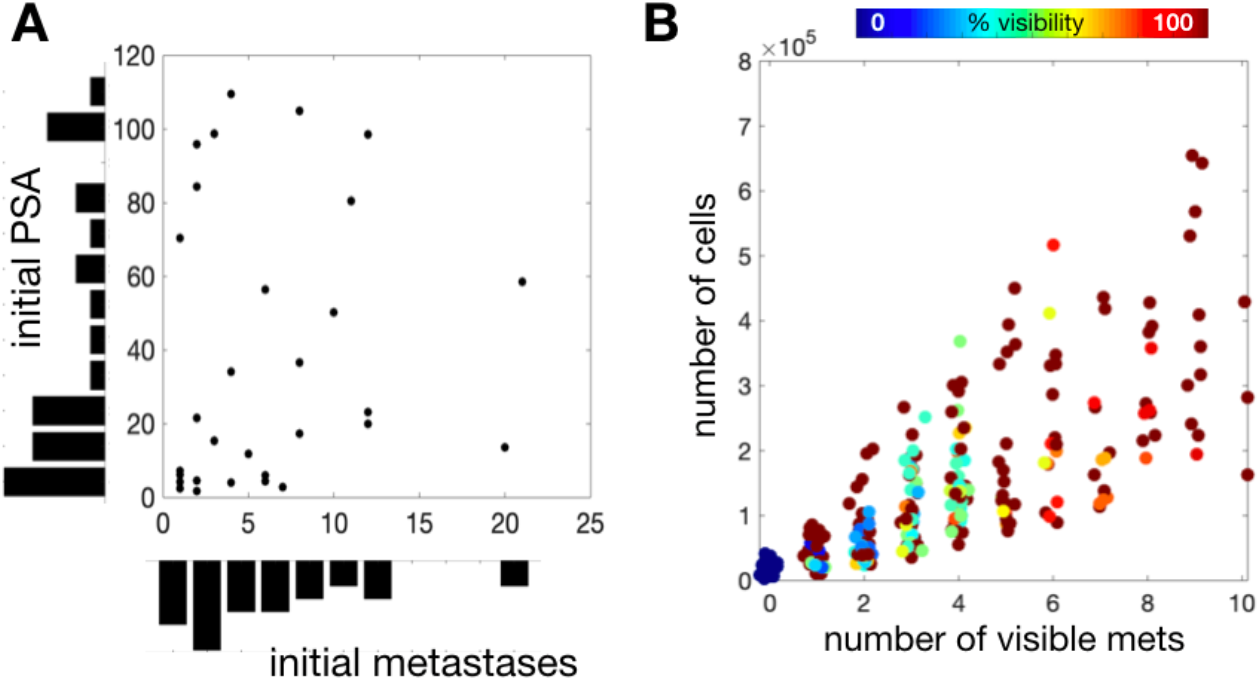
Burden vs. number of visible metastases for A) patients in the mCRPC the adaptive therapy trial (n=16) combined with the historical cohort (n=15) and B) the cohort of 1000 model simulations from Fig. 2E-F. While the patient data shows no apparent correlation between burden (PSA) and initial number of metastases, the simulated cohort does not have data points in the high burden, low metastases region.

**Figure S2.**
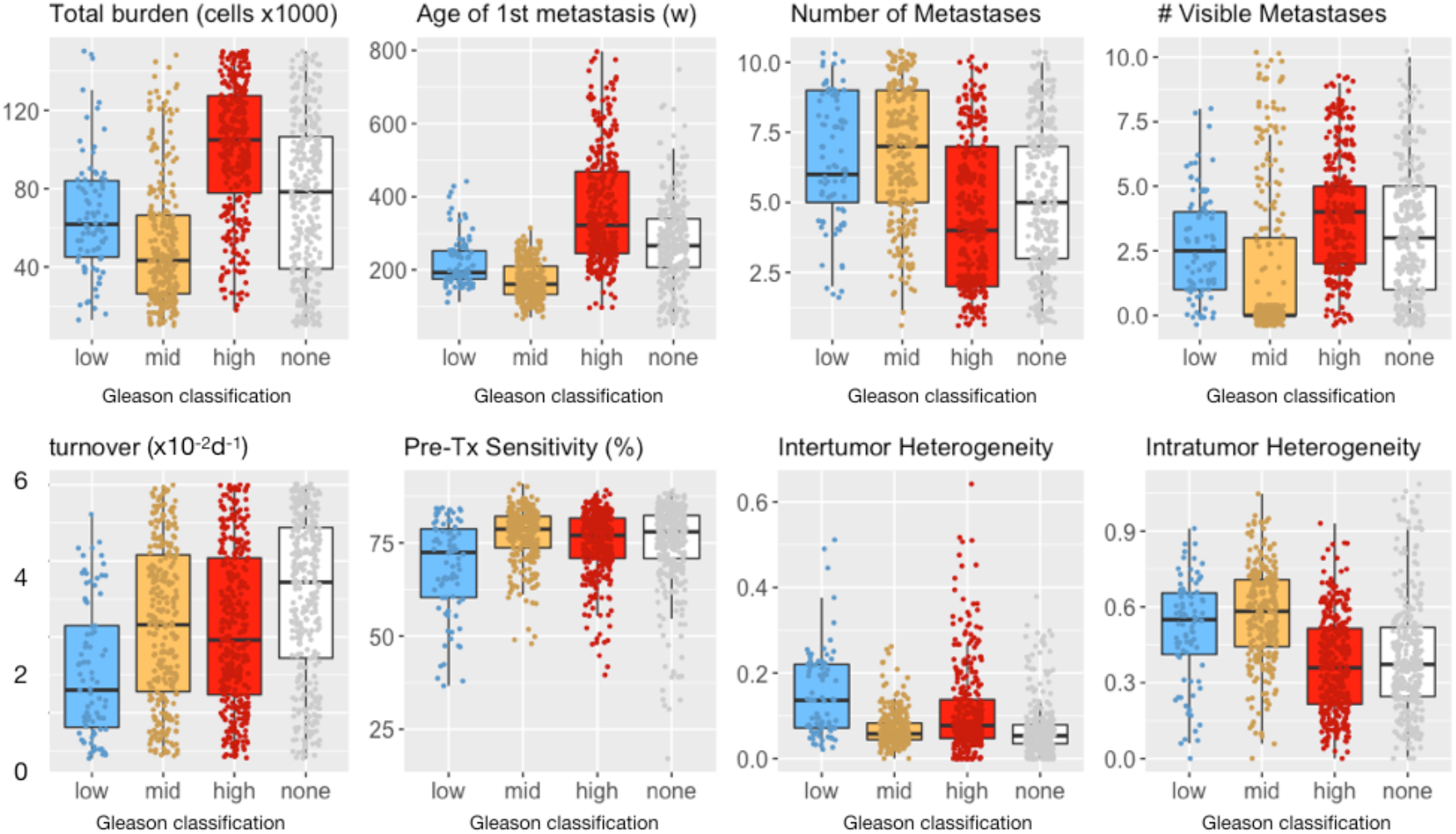
Comparison of the classified Gleason groups for the simulated data over different metrics. The classification is determined using the algorithm in the Supplemental Methods, which is based on data separated into Gleason Score (low=6-7, mid=8, and high=9-10). The category “none” did not fit the criteria to fit into any Gleason classification. The percent of samples in each group are 8.2%, 26.5%, 33.3%, and 32% in the low, mid, high, and none Gleason groups, respectively.

**Figure S3.**
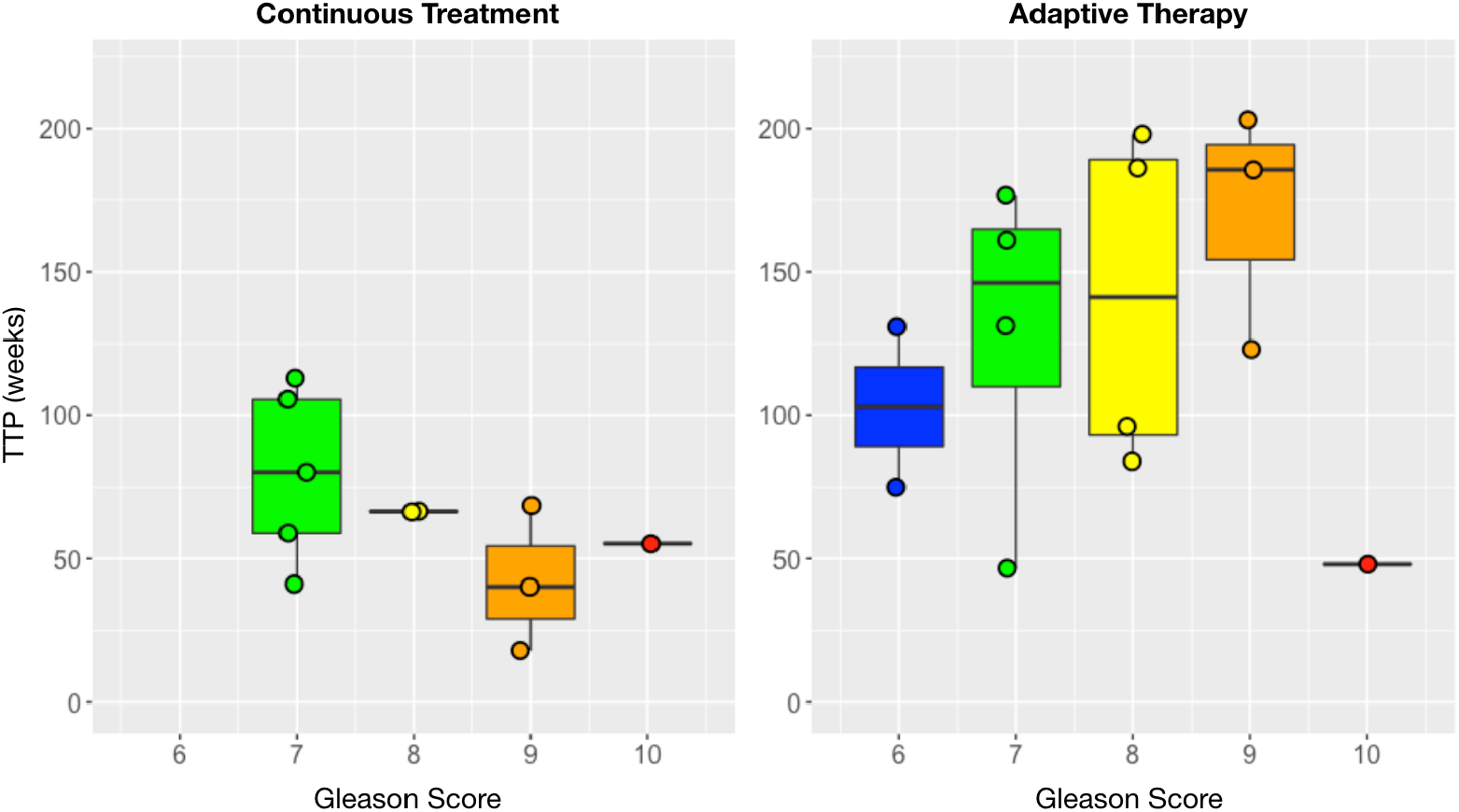
Comparison of time to progression (TTP) over different Gleason scores for continuous treatment (left) and adaptive therapy (right). A historical cohort is used for the continuous treatment patient data (n=12), and the mCRPC trial is used for the adaptive therapy patient data (n=15).

## Bibliography

[1] H. Dillekås, M. S. Rogers, and O. Straume, “Are 90% of deaths from cancer caused by metastases?,” Cancer Med-us, vol. 8, no. 12, pp. 5574–5576, 2019, doi: 10.1002/cam4.2474.

[2] D. S. Park et al., “The Goldilocks Window of Personalized Chemotherapy: Getting the Immune Response Just Right,” Cancer Res, vol. 79, no. 20, pp. 5302–5315, 2019, doi: 10.1158/0008-5472.can-18-3712.

[3] J. A. Scarborough, M. C. Tom, M. W. Kattan, and J. G. Scott, “Revisiting a Null Hypothesis: Exploring the Parameters of Oligometastasis Treatment,” Int J Radiat Oncol Biology Phys, vol. 110, no. 2, pp. 371–381, 2021, doi: 10.1016/j.ijrobp.2020.12.044.

[4] E. D. Crawford, C. L. Bennett, G. L. Andriole, M. B. Garnick, and D. P. Petrylak, “The utility of prostate-specific antigen in the management of advanced prostate cancer,” Bju Int, vol. 112, no. 5, pp. 548–560, 2013, doi: 10.1111/bju.12061.

[5] M. Hussain et al., “Absolute Prostate-Specific Antigen Value After Androgen Deprivation Is a Strong Independent Predictor of Survival in New Metastatic Prostate Cancer: Data From Southwest Oncology Group Trial 9346 (INT-0162),” J Clin Oncol, vol. 24, no. 24, pp. 3984–3990, 2006, doi: 10.1200/jco.2006.06.4246.

[6] P. M. Enriquez-Navas et al., “Exploiting evolutionary principles to prolong tumor control in preclinical models of breast cancer,” Sci Transl Med, vol. 8, no. 327, pp. 327ra24–327ra24, 2016, doi: 10.1126/scitranslmed.aad7842.

[7] R. A. Gatenby, A. S. Silva, R. J. Gillies, and B. R. Frieden, “Adaptive Therapy,” Cancer Res, vol. 69, no. 11, pp. 4894–4903, 2009, doi: 10.1158/0008-5472.can-08-3658.

[8] I. Smalley et al., “Leveraging transcriptional dynamics to improve BRAF inhibitor responses in melanoma,” Ebiomedicine, vol. 48, pp. 178–190, 2019, doi: 10.1016/j.ebiom.2019.09.023.

[9] J. Zhang, J. J. Cunningham, J. S. Brown, and R. A. Gatenby, “Integrating evolutionary dynamics into treatment of metastatic castrate-resistant prostate cancer,” Nat Commun, vol. 8, no. 1, p. 1816, 2017, doi: 10.1038/s41467-017-01968-5.

[10] Z. Kratiras, C. Konstantinidis, and K. Skriapas, “A review of continuous vs intermittent androgen deprivation therapy: Redefining the gold standard in the treatment of advanced prostate cancer. Myths, facts and new data on a ‘perpetual dispute,’” Int Braz J Urol, vol. 40, no. 1, pp. 3–15, 2013, doi: 10.1590/s1677-5538.ibju.2014.01.02.

[11] S. B. Strum, M. C. Scholz, and J. E. McDermed, “Intermittent Androgen Deprivation in Prostate Cancer Patients: Factors Predictive of Prolonged Time Off Therapy,” Oncol, vol. 5, no. 1, pp. 45–52, 2000, doi: 10.1634/theoncologist.5-1-45.

[12] J. M. Crook et al., “Intermittent Androgen Suppression for Rising PSA Level After Radiotherapy,” Obstet Gynecol Surv, vol. 68, no. 1, pp. 34–35, 2013, doi: 10.1097/01.ogx.0000426493.20419.c0.

[13] S. Magnan et al., “Intermittent vs Continuous Androgen Deprivation Therapy for Prostate Cancer: A Systematic Review and Meta-analysis,” Jama Oncol, vol. 1, no. 9, pp. 1–10, 2015, doi: 10.1001/jamaoncol.2015.2895.

[14] J. West, Y. Ma, and P. K. Newton, “Capitalizing on competition: An evolutionary model of competitive release in metastatic castration resistant prostate cancer treatment,” J Theor Biol, vol. 455, pp. 249–260, 2018, doi: 10.1016/j.jtbi.2018.07.028.

[15] L. You et al., “Spatial vs. non-spatial eco-evolutionary dynamics in a tumor growth model,” J Theor Biol, vol. 435, pp. 78–97, 2017, doi: 10.1016/j.jtbi.2017.08.022.

[16] J. J. Cunningham, J. S. Brown, R. A. Gatenby, and K. Staňková, “Optimal control to develop therapeutic strategies for metastatic castrate resistant prostate cancer,” J Theor Biol, vol. 459, pp. 67–78, 2018, doi: 10.1016/j.jtbi.2018.09.022.

[17] J. B. West, M. N. Dinh, J. S. Brown, J. Zhang, A. R. Anderson, and R. A. Gatenby, “Multidrug Cancer Therapy in Metastatic Castrate-Resistant Prostate Cancer: An Evolution-Based Strategy,” Clin Cancer Res, vol. 25, no. 14, pp. 4413–4421, 2019, doi: 10.1158/1078-0432.ccr-19-0006.

[18] M. A. R. Strobl, J. Gallaher, J. West, M. Robertson-Tessi, P. K. Maini, and A. R. A. Anderson, “Spatial structure impacts adaptive therapy by shaping intra-tumoral competition,” Commun Medicine, vol. 2, no. 1, p. 46, 2022, doi: 10.1038/s43856-022-00110-x.

[19] M. A. R. Strobl et al., “Turnover Modulates the Need for a Cost of Resistance in Adaptive Therapy,” Cancer Res, vol. 81, no. 4, pp. 1135–1147, 2021, doi: 10.1158/0008-5472.can-20-0806.

[20] E. Hansen, R. J. Woods, and A. F. Read, “How to Use a Chemotherapeutic Agent When Resistance to It Threatens the Patient,” Plos Biol, vol. 15, no. 2, p. e2001110, 2017, doi: 10.1371/journal.pbio.2001110.

[21] J. Zhang, J. Cunningham, J. Brown, and R. Gatenby, “Evolution-based mathematical models significantly prolong response to abiraterone in metastatic castrate-resistant prostate cancer and identify strategies to further improve outcomes,” Elife, vol. 11, p. e76284, 2022, doi: 10.7554/elife.76284.

[22] E. Kim, J. S. Brown, Z. Eroglu, and A. R. A. Anderson, “Adaptive Therapy for Metastatic Melanoma: Predictions from Patient Calibrated Mathematical Models,” Cancers, vol. 13, no. 4, p. 823, 2021, doi: 10.3390/cancers13040823.

[23] Y. Viossat and R. Noble, “A theoretical analysis of tumour containment,” Nat Ecol Evol, vol. 5, no. 6, pp. 826–835, 2021, doi: 10.1038/s41559-021-01428-w.

[24] R. Brady et al., “Prostate-Specific Antigen Dynamics Predict Individual Responses to Intermittent Androgen Deprivation,” Biorxiv, p. 624866, 2019, doi: 10.1101/624866.

[25] E. Hansen and A. F. Read, “Modifying Adaptive Therapy to Enhance Competitive Suppression,” Cancers, vol. 12, no. 12, p. 3556, 2020, doi: 10.3390/cancers12123556.

[26] A. S. Silva, Y. Kam, Z. P. Khin, S. E. Minton, R. J. Gillies, and R. A. Gatenby, “Evolutionary Approaches to Prolong Progression-Free Survival in Breast Cancer,” Cancer Res, vol. 72, no. 24, pp. 6362–6370, 2012, doi: 10.1158/0008-5472.can-12-2235.

[27] E. Ya. Tyuryumina and A. A. Neznanov, “Consolidated mathematical growth model of the primary tumor and secondary distant metastases of breast cancer (CoMPaS),” Plos One, vol. 13, no. 7, p. e0200148, 2018, doi: 10.1371/journal.pone.0200148.

[28] J. Gallaher, A. Babu, S. Plevritis, and A. R. A. Anderson, “Bridging Population and Tissue Scale Tumor Dynamics: A New Paradigm for Understanding Differences in Tumor Growth and Metastatic Disease,” Cancer Res, vol. 74, no. 2, pp. 426–435, 2014, doi: 10.1158/0008-5472.can-13-0759.

[29] S. Avanzini and T. Antal, “Cancer recurrence times from a branching process model,” Plos Comput Biol, vol. 15, no. 11, p. e1007423, 2019, doi: 10.1371/journal.pcbi.1007423.

[30] M. W. Retsky et al., “Computer simulation of a breast cancer metastasis model,” Breast Cancer Res Tr, vol. 45, no. 2, pp. 193–202, 1997, doi: 10.1023/a:1005849301420.

[31] K. Iwata, K. Kawasaki, and N. Shigesada, “A Dynamical Model for the Growth and Size Distribution of Multiple Metastatic Tumors,” J Theor Biol, vol. 203, no. 2, pp. 177–186, 2000, doi: 10.1006/jtbi.2000.1075.

[32] S. Benzekry and P. Hahnfeldt, “Maximum tolerated dose versus metronomic scheduling in the treatment of metastatic cancers,” J Theor Biol, vol. 335, pp. 235–244, 2013, doi: 10.1016/j.jtbi.2013.06.036.

[33] F. A. Coumans, S. Siesling, and L. W. Terstappen, “Detection of cancer before distant metastasis,” Bmc Cancer, vol. 13, no. 1, pp. 283–283, 2013, doi: 10.1186/1471-2407-13-283.

[34] L. C. Franssen, T. Lorenzi, A. E. F. Burgess, and M. A. J. Chaplain, “A Mathematical Framework for Modelling the Metastatic Spread of Cancer,” B Math Biol, vol. 81, no. 6, pp. 1965–2010, 2019, doi: 10.1007/s11538-019-00597-x.

[35] E. Szczurek, T. Krüger, B. Klink, and N. Beerenwinkel, “A mathematical model of the metastatic bottleneck predicts patient outcome and response to cancer treatment,” Plos Comput Biol, vol. 16, no. 10, p. e1008056, 2020, doi: 10.1371/journal.pcbi.1008056.

[36] A. Heyde, J. G. Reiter, K. Naxerova, and M. A. Nowak, “Consecutive seeding and transfer of genetic diversity in metastasis,” P Natl Acad Sci Usa, vol. 116, no. 28, pp. 14129–14137, 2019, doi: 10.1073/pnas.1819408116.

[37] L. A. Liotta, G. M. Saidel, and J. Kleinerman, “Stochastic Model of Metastases Formation,” Biometrics, 1976.

[38] L. A. Liotta, C. Delisi, G. Saidel, and J. Kleinerman, “Micrometastases formation: A probabilistic model,” Cancer Lett, vol. 3, no. 3–4, pp. 203–208, 1977, doi: 10.1016/s0304-3835(77)95675-0.

[39] P. Gerlee and M. Johansson, “Inferring rates of metastatic dissemination using stochastic network models,” Plos Comput Biol, vol. 15, no. 4, p. e1006868, 2019, doi: 10.1371/journal.pcbi.1006868.

[40] A. Rhodes and T. Hillen, “A mathematical model for the immune-mediated theory of metastasis,” J Theor Biol, vol. 482, p. 109999, 2019, doi: 10.1016/j.jtbi.2019.109999.

[41] J. G. Scott, D. Basanta, A. R. A. Anderson, and P. Gerlee, “A mathematical model of tumour self-seeding reveals secondary metastatic deposits as drivers of primary tumour growth,” J Roy Soc Interface, vol. 10, no. 82, p. 20130011, 2013, doi: 10.1098/rsif.2013.0011.

[42] J. Zhang, J. J. Cunningham, J. S. Brown, and R. A. Gatenby, “Response to Mistry,” Nat Commun, vol. 12, no. 1, p. 329, 2021, doi: 10.1038/s41467-020-20175-3.

[43] J. Zhang, J. J. Cunningham, J. S. Brown, and R. A. Gatenby, “Multidisciplinary analysis of evolution based Abiraterone treatment for metastatic castrate resistant prostate cancer,” Medrxiv, p. 2021.11.30.21267059, 2021, doi: 10.1101/2021.11.30.21267059.

[44] J. A. Gallaher, P. M. Enriquez-Navas, K. A. Luddy, R. A. Gatenby, and A. R. A. Anderson, “Spatial Heterogeneity and Evolutionary DynamicsModulate Time to Recurrence in Continuous andAdaptive Cancer Therapies,” Cancer Res, vol. 78, no. 8, p. canres.2649.2017, 2018, doi: 10.1158/0008-5472.can-17-2649.

[45] P. M. Pierorazio, P. C. Walsh, A. W. Partin, and J. I. Epstein, “Prognostic Gleason grade grouping: data based on the modified Gleason scoring system,” Bju Int, vol. 111, no. 5, pp. 753–760, 2013, doi: 10.1111/j.1464-410x.2012.11611.x.

[46] Z. Frankenstein et al., “Stromal reactivity differentially drives tumour cell evolution and prostate cancer progression,” Nat Ecol Evol, vol. 4, no. 6, pp. 870–884, 2020, doi: 10.1038/s41559-020-1157-y.

[47] K. W. Hunter, R. Amin, S. Deasy, N.-H. Ha, and L. Wakefield, “Genetic insights into the morass of metastatic heterogeneity,” Nat Rev Cancer, vol. 18, no. 4, pp. 211–223, 2018, doi: 10.1038/nrc.2017.126.

[48] C. A. Klein, “Parallel progression of primary tumours and metastases,” Nat Rev Cancer, vol. 9, no. 4, pp. 302–312, 2009, doi: 10.1038/nrc2627.

[49] Z.-M. Zhao et al., “Early and multiple origins of metastatic lineages within primary tumors,” Proc National Acad Sci, vol. 113, no. 8, pp. 2140–2145, 2016, doi: 10.1073/pnas.1525677113.

[50] G. Gundem et al., “The Evolutionary History of Lethal Metastatic Prostate Cancer,” Nature, vol. 520, no. 7547, pp. 353–357, 2015, doi: 10.1038/nature14347.

[51] A. Araujo, L. M. Cook, C. C. Lynch, and D. Basanta, “Size Matters: Metastatic Cluster Size and Stromal Recruitment in the Establishment of Successful Prostate Cancer to Bone Metastases,” B Math Biol, vol. 80, no. 5, pp. 1046–1058, 2018, doi: 10.1007/s11538-018-0416-4.

[52] Z. Hu, Z. Li, Z. Ma, and C. Curtis, “Multi-cancer analysis of clonality and the timing of systemic spread in paired primary tumors and metastases,” Nat Genet, vol. 52, no. 7, pp. 701–708, 2020, doi: 10.1038/s41588-020-0628-z.

[53] C. A. Klein and D. Hölzel, “Systemic Cancer Progression and Tumor Dormancy: Mathematical Models Meet Single Cell Genomics,” Cell Cycle, vol. 5, no. 16, pp. 1788–1798, 2006, doi: 10.4161/cc.5.16.3097.

[54] L. Willis et al., “Breast Cancer Dormancy Can Be Maintained by Small Numbers of Micrometastases,” Cancer Res, vol. 70, no. 11, pp. 4310–4317, 2010, doi: 10.1158/0008-5472.can-09-3144.

[55] K. Midde, N. Sun, C. Rohena, L. Joosen, H. Dhillon, and P. Ghosh, “Single-Cell Imaging of Metastatic Potential of Cancer Cells,” Iscience, vol. 10, pp. 53–65, 2018, doi: 10.1016/j.isci.2018.11.022.

[56] D. Merino et al., “Barcoding reveals complex clonal behavior in patient-derived xenografts of metastatic triple negative breast cancer,” Nat Commun, vol. 10, no. 1, p. 766, 2019, doi: 10.1038/s41467-019-08595-2.

[57] J. Pérez-Velázquez and K. A. Rejniak, “Drug-Induced Resistance in Micrometastases: Analysis of Spatio-Temporal Cell Lineages,” Front Physiol, vol. 11, p. 319, 2020, doi: 10.3389/fphys.2020.00319.

[58] J. P. Sleeman et al., “Concepts of metastasis in flux: The stromal progression model,” Semin Cancer Biol, vol. 22, no. 3, pp. 174–186, 2012, doi: 10.1016/j.semcancer.2012.02.007.

